# An arrayed CRISPR/Cas9 screen identifies mTORC1 as a regulator of lipid droplet accumulation in APOE E3 and APOE KO iPSC-derived microglia

**DOI:** 10.1101/2024.03.16.585330

**Authors:** Sonja Meier, Anne Sofie Gry Larsen, Florian Wanke, Nicolas Mercado, Arianna Mei, Livia Takacs, Eva Suszanna Mracsko, Ludovic Collin, Martin Kampmann, Filip Roudnicky, Ravi Jagasia

## Abstract

Variants of the Apolipoprotein E (APOE) gene, particularly the E4 allele, are significantly associated with an increased risk of Alzheimer’s Disease and have been implicated in neuroinflammatory processes due to disrupted lipid metabolism. Lipid alterations can manifest in glial cells as an excessive buildup of lipids, potentially contributing to neuroinflammation. In this study, we observed a heightened lipid load in APOE-deficient human induced pluripotent stem cell (iPSC)-derived microglia relative to cells with other APOE isoforms. To explore the mechanisms governing lipid handling within microglia, we established a technique for the nucleofection of CRISPR/Cas9 ribonucleoprotein complexes into iPSC-derived myeloid cells. Utilizing this method, we performed a targeted screen to identify key upstream modifiers in lipid droplet formation. Our findings highlight the mammalian target of rapamycin complex 1 (mTORC1) signaling pathway as a pivotal influence on lipid storage in microglia with both APOE3 and APOE knockout genotypes, underscoring its role in lipid dysregulation associated with Alzheimer’s Disease and neuroinflammation.

## Introduction

Pathologically, Alzheimer’s Disease (AD) is characterized not only by amyloid plaque and tau tangles but also lipid-laden glial cells^1^. Genome-wide association studies (GWAS) of late onset AD (LOAD) have revealed genes enriched in both lipid metabolism and innate immunity^2^. *APOE* is a major LOAD risk gene linked to lipid metabolism and is highly upregulated in human microglia in AD^3,4^. Notably, both postmortem and human iPSC-derived microglia with *APOE* risk variant ε4 have more lipid droplets, and this accumulation impairs microglia homeostatic functions and leads to a shift towards a proinflammatory state^5^.

Despite the growing genetic evidence that glial lipid dysfunction plays a role in LOAD pathophysiology in part via APOE, the mechanism remains poorly understood. Studies examining lipid dysregulation in microglia have focused largely on measuring the relationship, not causality, between the amount of lipids, and the expression of lipid and inflammatory related genes. This can, in part, be attributed to the historical lack of robust methods for genetic manipulation of myeloid cells including microglia^6^. So far, CRISPR-based microglia studies have involved genetic manipulations at the pluripotent stem cell stage^7^, which poses a challenge for both arrayed screens with high-content readouts, and also to uncouple developmental vs. functional effects in mature cells. To overcome this challenge, we developed an efficient method for nucleofection of iPSC-derived macrophages and microglia and conducted an arrayed screen to determine factors that positively and negatively regulate lipid accumulation in APOE3 and APOE KO iPSC-derived microglia.

## Results

### APOE regulates lipid droplet accumulation in iPSC-derived microglia

To establish a model that can be used for arrayed genetic screening to find genes that regulate lipid droplet formation, we characterized the APOE lipid droplet phenotype in microglia and macrophages derived from isogenic iPSCs harboring different APOE alleles, or an APOE KO (Figure S1B). Myeloid factories were generated as previously described^8,9^ and macrophage precursors (‘preMacs’) were harvested continuously from week 6 to 12 (as long as they passed quality control; Figure S1A) for further differentiation. To image lipid droplets, we took advantage of Nile Reds (NR) ability to excite at different wavelengths depending on its binding to polar or non-polar lipids^10^, allowing visualization of both cell membrane and lipid droplets (Figure 1). APOE KO microglia had a significantly increased number of lipid droplets, based on both ‘spots’ per cell area and NR intensity per cell area, compared to APOE3 (Figure 1A, B). APOE4 microglia, but not APOE2, showed a trend towards increased lipid content that was non-significant (Figure 1A, B). This effect was not due to a difference in APOE expression (Figure S1C). Interestingly, APOE KO resulted in a significant increase in cell size, with a ∼40% increased cell area vs APOE3. Consistently, iPSC-derived macrophages showed a significant increase in the number of lipid droplets in both APOE KO and E4 compared to E3 (Figure S1D, E). Since the most prominent effect was observed for APOE KO, we decided to use APOE3 and APOE KO lines to screen for lipid modulators in iPSC-derived microglia.

**Figure 1:**
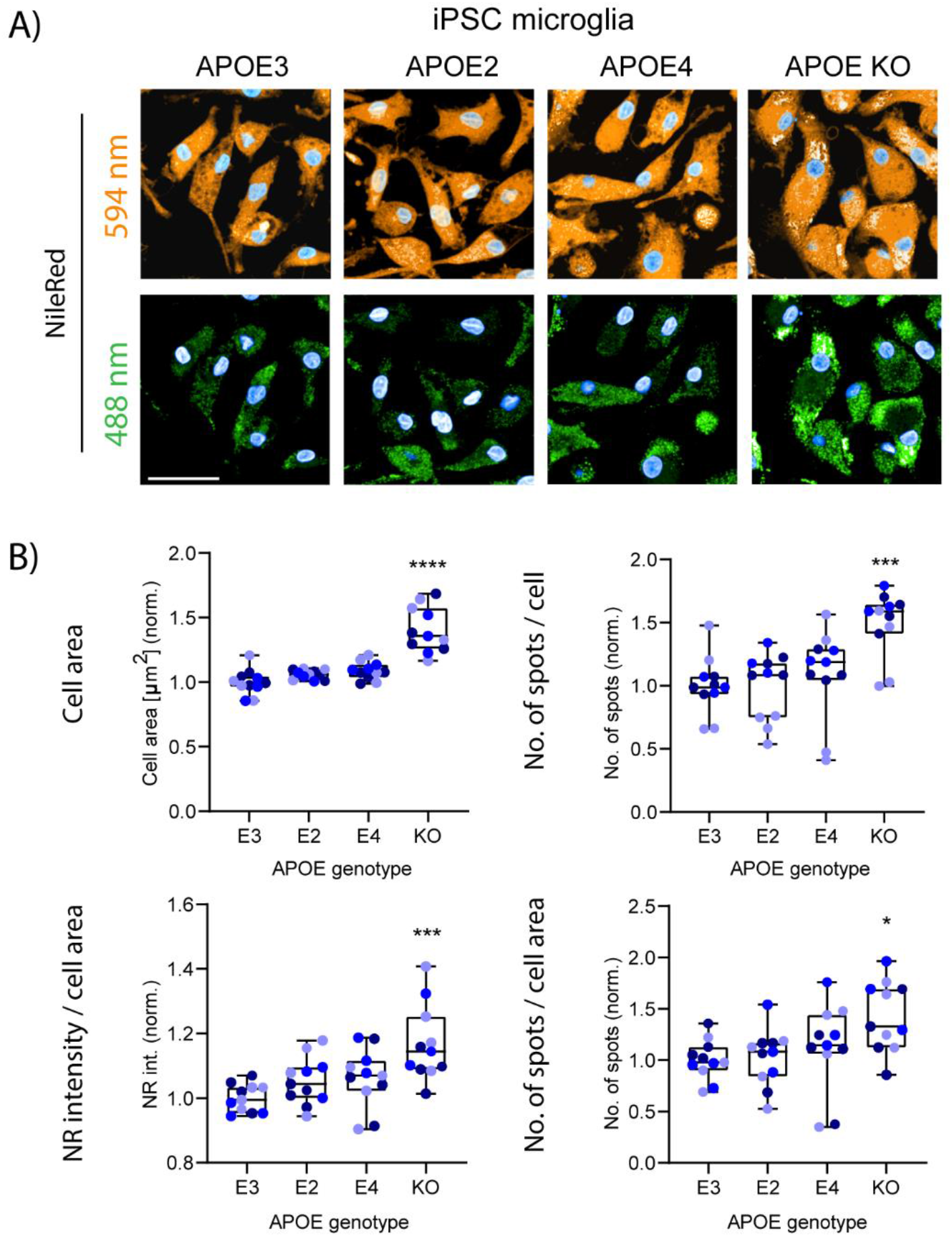
APOE regulates intracellular lipid accumulation and cell size in iPSC-derived microglia. **A)** Representative images of iPSC-microglia stained with Nile red (NR) and imaged at 488 nm (non-polar lipids) and 594 nm (polar lipids). Scale bar: 50 µm. **B)** NR intensity and number of spots (488 nm) normalized to cell area or per quantified number of cells (594 nm). Different shades of blue indicate 3 independent biological experiments of minimum 3 technical replicates. Data are normalized to APOE3 for each experiment. One-way ANOVA for paired values across experiments was performed, followed by Dunett’s multiple comparison correction.

### Optimization of CRISPR/Cas9 nucleofection for iPSC-derived myeloid cells

To allow efficient gene knockout in iPSC-derived myeloid cells without compromising the cells’ antiviral properties, we optimized a protocol for delivery of the CRISPR/Cas9 machinery via nucleofection in microglia, macrophages and preMacs. Initial conditions were based on a previously published protocol for PBMCs^11^ using CD81 sgRNA for optimization. As the base media is different for different iPSC-derived microglia and macrophage differentiation methods^12^, we first tested the effect of different base media including X-VIVO15, ImmunoCult-SF, RPMI 1640 + 10% FCS, and DMEM/F12 + N2 on cell viability post nucleofection. Both DMEM/F12 and RPMI 1640 culture media yielded >90% viability of iPSC-derived macrophages 5 days after nucleofection as determined by flow cytometry. In contrast, cell survival in X-VIVO15 and ImmunoCult XF was <20% and <50%, respectively (Figure S2A). In more than 80% of surviving cells, CD81 was successfully knocked out for all conditions (Figure S2A). We next investigated whether KO efficiency and/or survival could be further improved by changing the pulse code settings. Testing 15 different programs, we confirmed CM-137 as the most robust condition with 70-80% survival and ∼80% CD81 KO (Figure S2B, C). Finally, we optimized the Cas9 : sgRNA ratio for RNP formation to 2 µg Cas9 : 25 nmol sgRNA (Figure S2D). To ensure that CRISPR/Cas9 KO by nucleofection would translate into a functional response, we used these optimized conditions to knockdown TLR4. As expected, secretion of TNFα and IL-6 in response to LPS treatment, but not poly-I:C, was impaired in the TLR4 KO compared to NTC (Figure S2E). Thus, we have successfully optimized a method to regulate myeloid function through nucleofection of the CRISPR/Cas9 machinery.

### CRISPR/Cas9 nucleofection can be used for gene knockout in both developing and mature iPSC-derived microglia and macrophages

To ensure that CRISPR/Cas9 nucleofection could also be employed to microglia, we tested our optimized method in both mature microglia (day 10 of differentiation) as well as preMacs that were subsequently differentiated into microglia or macrophages. Using CD81 sgRNA together with Cas9, we observed an >80% CD81 gene KO for all conditions (Figure 2A). For all subsequent experiments, we used iPSC-derived microglia nucleofected at preMac stage. We next asked whether nucleofection would affect microglial activation. Microglia were stained using antibodies against IBA1, TMEM119, or P2Y12 after nucleofection with Cas9 and NTC, CD81, or OR52A1 (an olfactory gene not expressed in microglia) and compared to non-nucleofected controls (‘Ctrl’; Figure 2B). Microglia of all conditions stained positive for IBA1, TMEM119, and the resting marker P2Y12 (Figure 2C). No obvious differences in cell roundness or cell size were detected compared to control, indicating that nucleofection did not shift microglia morphologically towards an activated state. To test whether nucleofection affected a classical microglial function, secretion of TNFα and IL-6 was assessed at baseline and after LPS treatment. Notably, nucleofected myeloid cells responded to LPS treatment, although with a slightly diminished response compared to non-nucleofected controls (Figure 2D). Since microglial activation can affect lipid metabolism, we further asked whether nucleofection would affect the lipid accumulation we observed in APOE KO microglia (Figure 1). Although NR intensity per cell area was increased across all APOE variants following nucleofection, the APOE-dependent lipid phenotype was maintained (Figure 2E, F).

**Figure 2:**
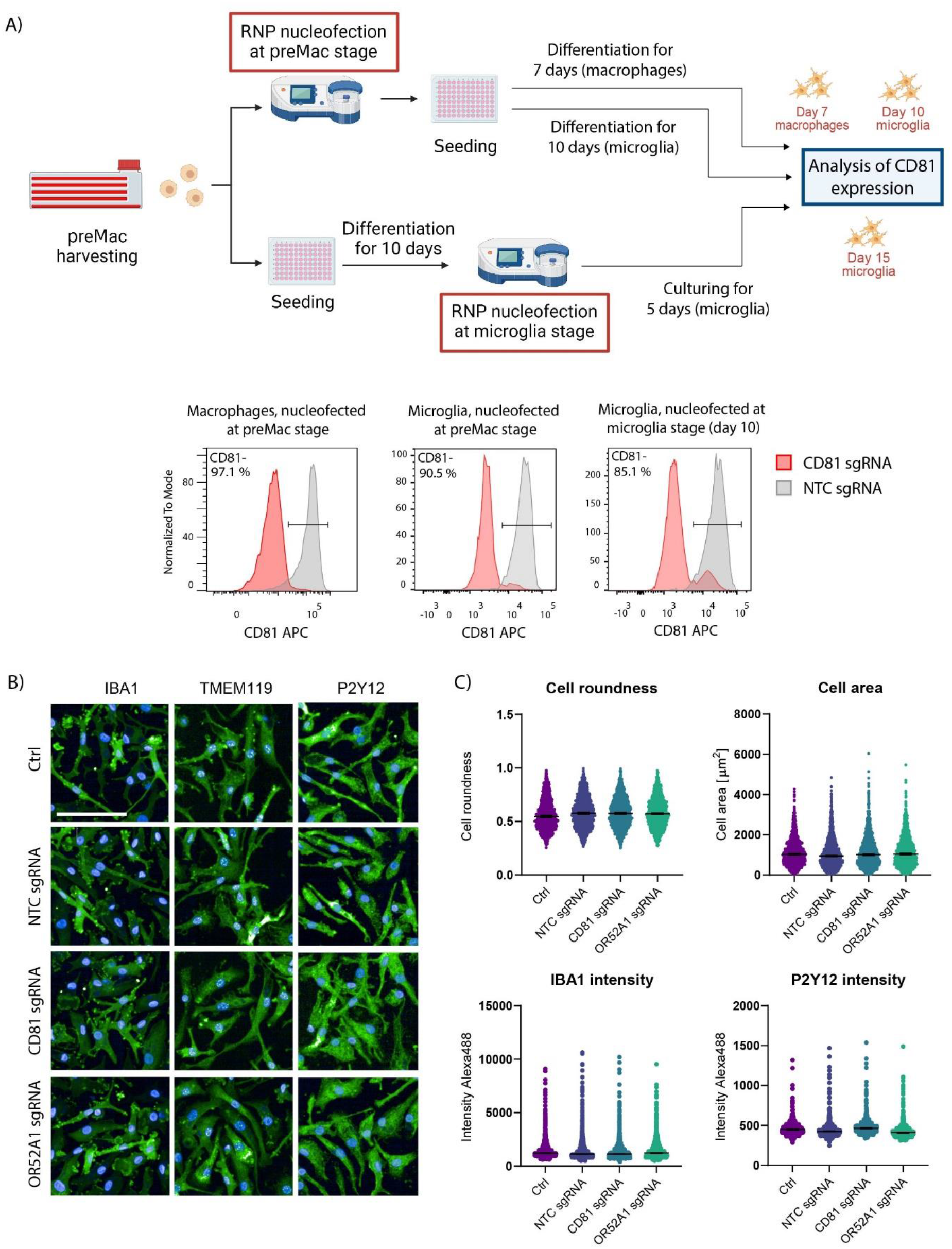
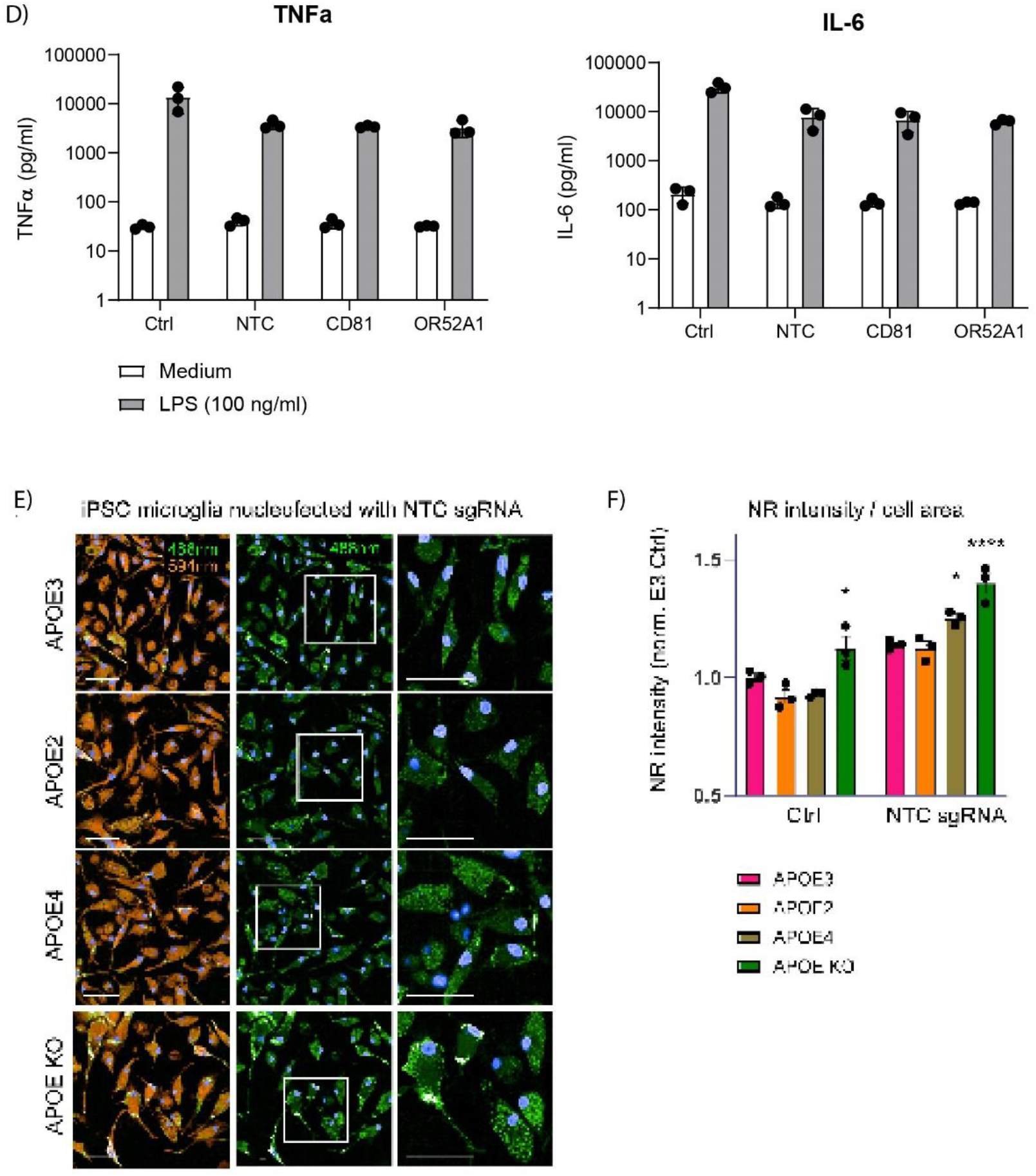
CRISPR/Cas9 RNP nucleofection can effectively edit iPSC-derived microglia, macrophages or preMacs without altering their function. **A)** Microglia and preMacs differentiated into macrophages or microglia were nucleofected with RNPs containing non-targeting control (NTC) or CD81 sgRNA. CD81 expression was analyzed after 7 days (preMacs to macrophages), or 10 days (preMacs to microglia) or 5 days (mature microglia; left to right) by flow cytometry. **B)** Representative images of microglia immunofluorescently stained using antibodies against IBA1, TMEM119, and P2Y12 after nucleofection at preMac stage with RNPs containing NTC, CD81, or OR52A1 sgRNA. Ctrl = non-nucleofected cells. Scale bar: 100 µm. **C)** Quantification of immunofluorescence single-cell data with population median and 95% confidence interval. **D)** Microglia treated with 100 ng/ml LPS for 6 hours before supernatant collection for Luminex cytokine measurement. Data from three independent experiments. **E)** Microglia nucleofected at preMac stage with Cas9 and NTC sgRNA and stained with NR. Scale bar: 50 µm. **F)** Quantification of NR intensity (488 nm) per cell area, normalized to APOE3 Ctrl. 2-way ANOVA followed by Dunnett post-hoc testing comparing all lines to APOE3 within each nucleofection group. Data representative for two independent experiments that were analyzed independently.

### Arrayed screen in iPSC-derived microglia reveals positive and negative regulators of lipid droplet formation

To conduct an arrayed gene KO screen using our nucleofection method, we compiled a focused sgRNA library containing 334 genes. In addition to neurodegeneration associated genes, we targeted genes involved in intracellular lipid homeostasis, including uptake, modification, storage, signaling, and efflux of lipids (Table S1). Three sgRNAs per target and per well were used and randomly distributed among four 96-well plates. Each plate also contained five different NTCs (Table S1), two positive controls to assess KO efficiency (CD81, CD14), three DNA cutting controls (targeting olfactory genes not expressed in myeloid cells), and two kill controls (essential genes deletion of which is predicted to be toxic; Table S1). Each sgRNA plate was used in three independent nucleofection reactions. After nucleofection, preMacs of each sgRNA plate were seeded into two replicate 96-well plates (Figure 3A). On day 10 of differentiation, KO efficiency was confirmed via flow cytometry of the cell surface markers (CD81 and CD14) and remaining wells were fixed and stained with NR for lipid droplet analysis. We noticed an inverse correlation between cell number and NR intensity per cell area, with fewer cells resulting in increased NR signal (Figure S3B). We concluded that in wells with reduced number of cells, increased lipid signal may not be directly linked to target gene function but could be a product of unspecific stress or increased phagocytosis of dead cells by surviving microglia. We therefore excluded genes for which cell number differed >20% from that of average NTC, resulting in 56 and 31 genes being excluded from APOE3 and APOE KO datasets, respectively (Figure S3C). Of the remaining genes, we defined hits as >15% increased or decreased NR intensity per cell area. Most hits increased lipid content in both APOE3 and APOE KO microglia, with 44 and 53 targets increasing, and only 3 and 5 targets decreasing lipids, respectively (Figure 3B, C). No significant effect on NR intensity was observed for DNA cutting controls (Figure S3D). Importantly, KO of Sterol-O-Acyl transferase 1 (SOAT1/ACAT1), an enzyme transforming free cholesterol into its CE storage form, strongly reduced lipid accumulation independent of APOE genotype, as seen previously for SOAT1 inhibitors^13,14^. We confirmed this effect in microglia derived from an independent iPSC line (BIONi010-C; *APOE3/4*) and compared it with a SOAT1 inhibitor. Both KO and pharmacological inhibition significantly decreased NR intensity per cell area, as expected (Figure 3D, E). Compared to the NR intensity readout, the number of lipid droplets was only meaningfully changed for a handful of targets (Figure S4A, B). Of note is that KO of another well-known regulator of lipid storage, Diacylglycerol O-acyltransferase 1 (DGAT1)^15-17^, significantly decreased the number of lipid droplets in APOE3, but not APOE KO microglia, which might suggest differences in lipids species accumulating in the two lines. The effect on lipid droplet number by DGAT1 KO or pharmacological inhibition was also independently confirmed in BIONi010-C-derived microglia (Figure 3D). Many hits increasing NR intensity were found to either localize to the lysosome, or be involved in lysosomal regulation, including LGMN, LAMP1, LAMP2, LIPA, CTSC, SCARB1, VPS35, VPS39, RAB7A, LAPTM4B, STARD3NL, ABCA2, MAP1LC3A, and TFEB, as well as components of the mTORC1 pathway.

**Figure 3:**
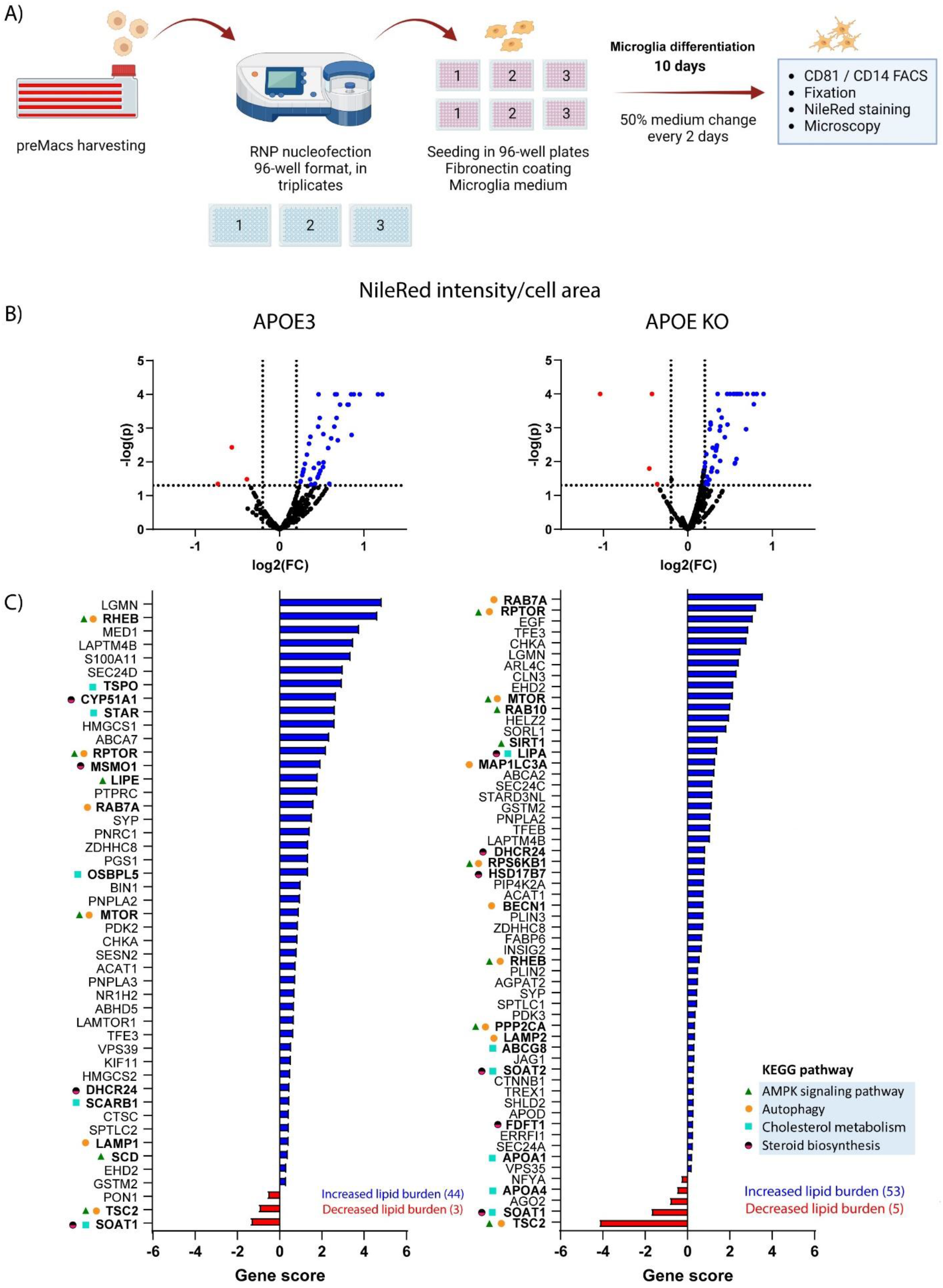
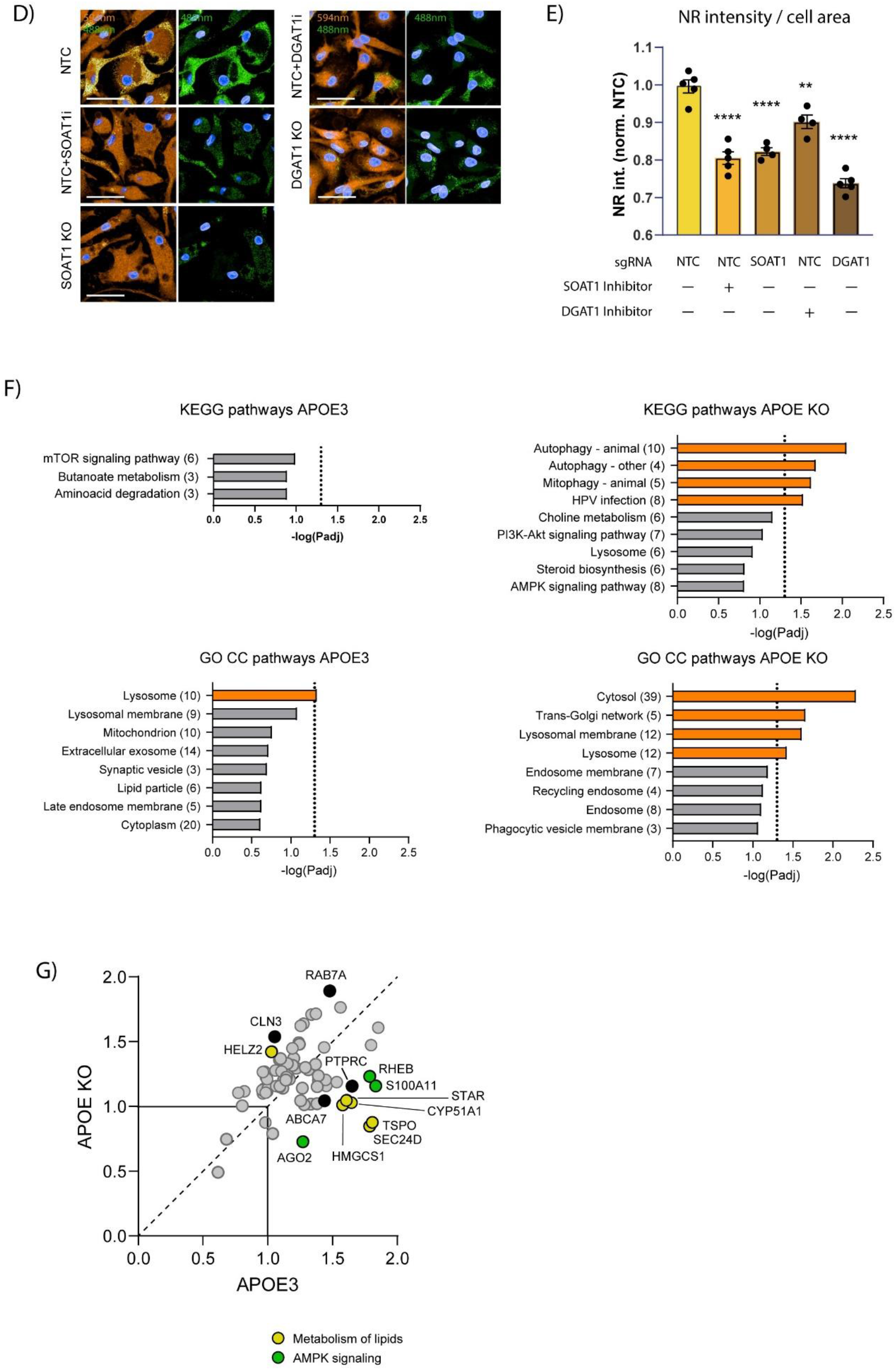
Arrayed lipid droplet screen reveals positive and negative regulators in APOE3 and APOE KO microglia. **A)** Schematic representation of the arrayed CRISPR/Cas9 screen workflow. **B)** Volcano plots indicating significant hits for APOE3 and APOE KO for Nile red (NR) intensity quantification. Red and blue dots indicate significant hits with an increase in NR intensity of at least 15%. Data averaged from three nucleofection experiments per gene. Every gene was analyzed for significance separately for each experiment with paired 1-way ANOVA and comparison to NTC on each plate. **C)** Gene scores for significant hits of APOE3 and APOE KO. Gene score = log2(FC) * -log(p). Blue bar color indicates increased NR intensity / cell area, red bar color indicates decreased intensity. **D, E)** Effect of SOAT1 KO and DGAT1 KO on NR intensity/cell area in BIONi010-C (APOE3/4) microglia. Inhibitor treatment: SOAT1 inhibitor (K604) 500 nM, or DGAT1 inhibitor (PF-04620110) 1 µM, for 2 days. 2-way ANOVA with Sidak post-hoc test. **F)** KEGG pathway and GO-term cellular component (GO CC) analysis using DAVID Bioinformatics Database (version 2021; threshold 2; EASE 0.25; Benjamini correction) of significant hits on the background of the genes and corresponding pathways represented in the focused library. **G)** Correlation plot of hits APOE3 vs. APOE KO for NR intensity. Only hits that did not reduce cell number in both lines were analyzed. 2-way ANOVA was used to determine significant differences between the lines (colored circles = significant, grey circles = non-significant).

### Comparison of gene KO effects on lipid content reveals key pathways

To better understand the pathways regulating lipid content, and their dependence on APOE, we performed KEGG pathway analysis (Figure S4C). First, we compared top pathways of the sgRNA library as well as the significant hits for both APOE3 and APOE KO with g-profiler (Figure S4D). The top four pathways represented in the library, steroid biosynthesis, autophagy, cholesterol metabolism, and AMPK signaling were enriched in both lines, although APOE KO scored higher on autophagy compared to APOE3 (10 vs 6 with 5 common genes; Figure S4D). A notable exception is the PPAR signaling pathway, which did not appear in the top 9 enriched pathways of either line albeit being the second most represented pathway in the library (Figure S4D, Table S3). Interestingly, enriched hits in cholesterol metabolism were largely non-overlapping between lines, except for SOAT1, potentially suggesting differences in lipid species accumulation. We next performed an enrichment analysis on the background of the complete gene list using DAVID Bioinformatics Database to correct for targeted library bias^18^. Although mTOR signaling scored highest for APOE3, no pathway was enriched significantly above background, while autophagy, mitophagy, and HPV infection pathways were significantly enriched in APOE KO microglia (Figure 3F).

Comparing individual hits between lines revealed that more targets increase lipid content in APOE3 microglia compared to APOE KO, likely due to a higher baseline lipid content in APOE KO (Figure 3G). Among targets increasing NR signal in APOE3 but not APOE KO microglia are TSPO and STAR, two outer mitochondrial proteins involved in shuttling cholesterol into mitochondria^19^ (Figure 3G). Notable similarities between APOE3 and APOE KO microglia were hits directly influencing the mTORC1 pathway.

### mTORC1 signaling is a crucial regulator of lipid droplet formation and cell size in iPSC-derived microglia

While KO of TSC1 did not change lipids significantly, TSC2 KO (and therefore mTORC1 activation) strongly reduced lipid content in both genotypes. Conversely, KO of genes positively regulating mTORC1 signaling increased lipids in both lines including RHEB, RPTOR and MTOR itself (Figure 3G). We chose the top positive and negative regulators from the mTOR pathway in APOE3 microglia, RHEB and TSC2, for KO validation in microglia derived from the independent BIONi010-C line (*APOE3/4*). We compared this to acute (24 h) and chronic (5 day) mTORC1 inhibition by rapamycin treatment. We observed increased mTORC1 activity, determined as phospho-S6 / total S6 level, in TSC2 KO cells (Figure 4D). 5-day rapamycin treatment caused a large decrease in phospho-S6 / total S6 while RHEB KO and 24 h rapamycin treatment had minor effects. The effects of TSC2 and RHEB KO on lipid droplet formation and cell area confirmed the findings from the CRISPR/Cas9 screen, with TSC2 and RHEB decreasing and increasing NR intensity per cell area as well as increasing and decreasing cell area, respectively (Figure 4C, E, F). Interestingly, the effect of RHEB KO was stronger than that observed for chronic rapamycin treatment despite the minor effect of RHEB KO on phospho-S6 / total S6. For TSC2 KO, both the decreased NR intensity and increased cell size could be reversed by rapamycin treatment (Figure 4E, F). Collectively, this data points to mTORC1 signaling playing an important role in regulating lipid storage in iPSC-derived microglia.

**Figure 4:**
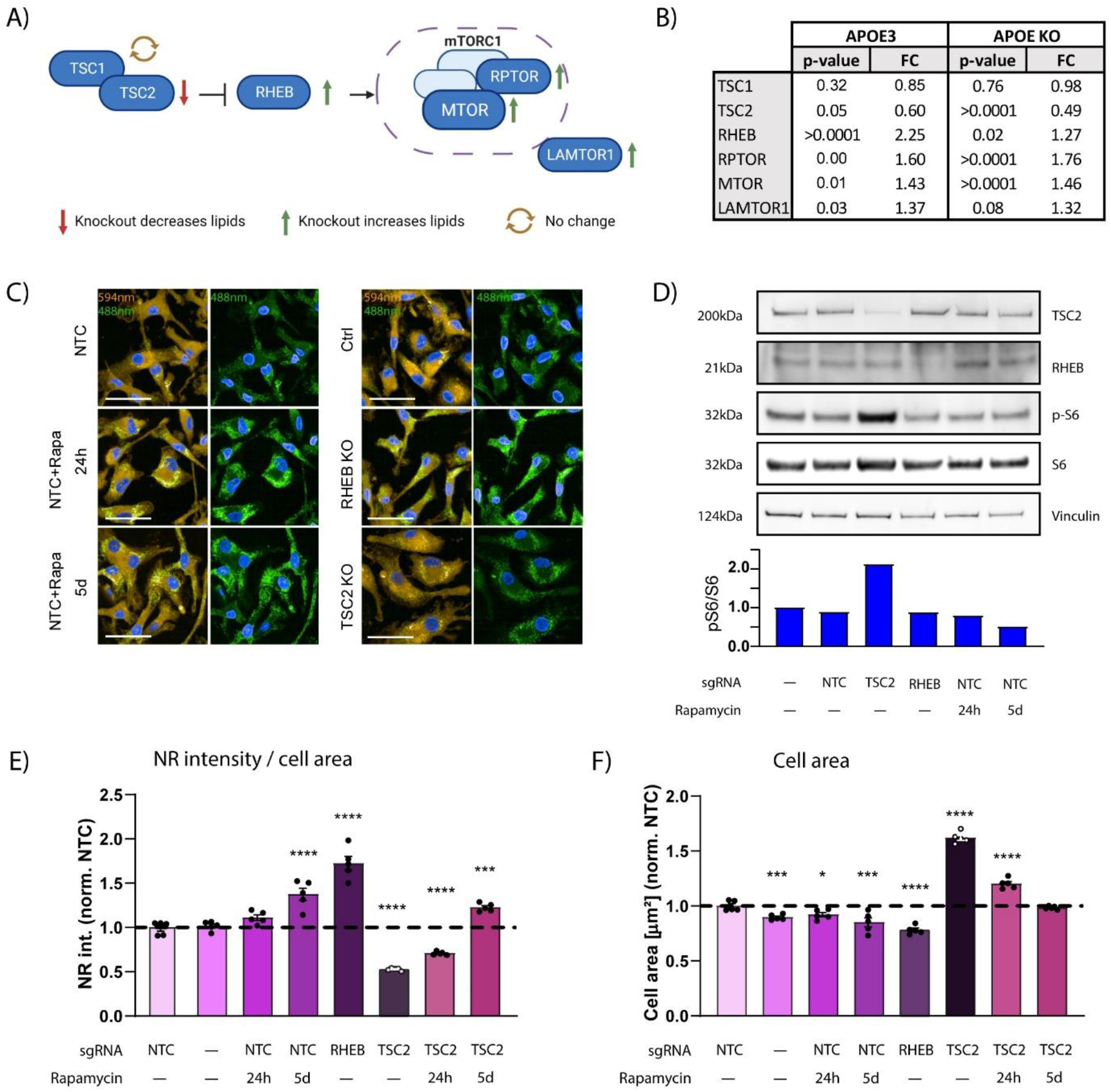
mTORC1 signaling regulates lipid droplet accumulation and cell size in iPSC-derived microglia. **A)** Schematic representation of the mTORC1 pathway and the effect of KO on lipid load as identified in the CRISPR/Cas9 arrayed screen. **B)** Report of fold change and p-values for mTORC1 signaling pathway components tested in the CRISPR/Cas9 arrayed screen. **C)** Validation of RHEB and TSC2 KO on lipid phenotype in microglia from BIONi010-C line measured by NR staining. Rapamycin treatment with 2 nM for 24 h or 5 days before analysis. (NTC-non-targeted control sgRNA) **D)** Western blot for TSC2, RHEB, p-S6 and S6 in BIONi010-C microglia. **E, F)** Quantification of NR intensity (488 nm) and cell area (594 nm) of images shown in C). Data representative of 2 independent experiments; analyzed using 2-way ANOVA.

## Discussion

Cholesterol dysfunction is a key component of AD pathophysiology contributing to microgliosis, a major hallmark of the disease^20-24^. Several LOAD genetic risk factors have important functions in microglial cholesterol metabolism^13,25,26^, yet a mechanistic understanding of the pathways involved in lipid dysfunction in the context of AD is largely lacking. The paucity of knowledge is in part due to challenges in genetic manipulations of microglia, which resist viral transduction via lentivirus or AAV^27^. To that end, we now present a method for delivering the CRISPR/Cas9 machinery into iPSC microglia through nucleofection of the RNP, allowing gene KO with >80% efficiency to rapidly and efficiently perform genetic loss-of-function studies. We tailored this approach to perform a targeted CRISPR/Cas9 KO screen to investigate genes involved in intracellular lipid handling in microglia in an APOE-dependent manner. We identify both known and novel genes modulating lipid droplets and demonstrate an unexpected role of mTORC1 activation in reducing lipid accumulation independent of APOE.

Most hits from the targeted lipid droplet screen in both APOE3 and APOE KO microglia resulted in an increase of non-polar lipid cellular content, whilst only few hits reduced lipid levels. The fact that we identified mostly positive regulators is in part due to an overrepresentation of genes involved in lipid handling in our 334 gene focused library. For example, it is well known that lysosomal dysfunction results in altered lipid metabolism and increased lipid droplet formation in conditions known as Lysosomal Storage Disorders^28,29^. This was captured in the screen, as many of the identified hits were either physically located to the lysosome, or indirectly involved in lysosomal biogenesis and function. For instance, *LIPA* encodes Lysosomal Acid Lipase (LAL) that hydrolyzes cholesteryl esters to release free cholesterol, therefore catalyzing the reverse reaction from SOAT1/ACAT1 and reducing lipid load^12^. Other hits included lysosomal enzymes not likely involved in lipid processing directly, such as the luminal cysteine proteases CTSC and LGMN, in the case of which loss-of-function may contribute first to lysosomal dysfunction via substrate build up and in lipid changes as a secondary effect^29^.

We observed a small net increase of non-polar lipid levels in microglia after RNP nucleofection, likely as result of cellular stress response or to phagocytosis of dead cells after nucleofection. Indeed, the majority of gene KO that inhibited microglial survival increased lipid levels. Beyond core lysosomal genes, biologically meaningful increases in lipids were seen in genes associated with mTORC1, AMPK and autophagy signaling pathways. Whilst there was a good overlap between hits in both APOE3 and APOE KO microglia, the effects were less pronounced in the latter, likely due to the increased baseline lipid levels. Interestingly however, APOE KO microglia appeared more sensitive to loss-of-function of autophagy and lipophagy-related genes. For example, deletion of MAP1LC3/LC3, which interacts with motifs of specific autophagy receptors and has been implicated in lipophagy-mediated lipid droplet catabolism in macrophages^20^, resulted in a significant increase in lipid content in APOE KO, but not APOE3 microglia. Deleting the crucial autophagy gene *Atg*5 in macrophages in an APOE KO mouse model of atherosclerosis led to an acceleration in the acquisition of the atherogenic phenotype, including increased inflammasome activation and macrophage recruitment to plaques^21^. In a different study of autophagy-deficient (Atg7-/-) microglia in mice, an increase in the amount of lipid droplets was found, which was further exacerbated by LPS treatment concomitantly with upregulation of APOE expression^22^. Collectively, this could suggest an axis whereby APOE may help clear lipids in the context of degradation defects, while APOE dysfunction leads to increased dependency on lipophagy.

Our screen also captured AD risk genes with both described and unknown roles in lipid metabolism, including ABCA7 (involved in cholesterol metabolism and amyloid processing^30^), BIN1 (linked to endolysosomal function^31^), SORL1 (a low-density lipoprotein receptor and involved in amyloid precursor protein trafficking^32^), NR1H2 (LXR-B; linked to neuro-inflammation^33^), and targets associated with the VPS35/retromer complex (involved in microglial phagocytosis, lysosomal transport and homeostasis^34,35^). While future work is required to delineate the precise pathomechanisms of these targets, this study represents the first hint that glial associated lipid pathology could be a contributing factor.

Both top negative and positive regulators of lipid load are functionally associated with mTORC1, a major metabolic pathway at the intersection of autophagy, lysosomal function, and lipid metabolism^36^. Activated mTORC1 is anchored to the outer lysosomal membrane and prevents autophagosome formation through inhibitory phosphorylation of TFEB and other lysosome-associated proteins^23,24^. mTORC1 also regulates lipid homeostasis via Sterol Regulatory Element-Binding Proteins (SREBPs), which are transcriptional master regulators of lipogenesis^25^. Therefore, it was paradoxical that both genetic and pharmacological inhibition of mTORC1 led to an increased lipid burden in our microglia. Interestingly, a recent study investigating mTORC1-controlled lipid accumulation in several cell lines found that de novo lipid synthesis upon mTOR activation mainly produced phospholipids for membrane growth, while mTOR inhibition led to a reversal and lysosome-dependent breakdown of phospholipids into triglycerides that are deposited in lipid droplets for energy storage^37^. Therefore, the lipids accumulating in mTORC1-suppressed microglia would be expected to primarily be triglyceride species, and fewer cholesteryl esters, as was found in hepatocytes of a *Rheb-/-* mouse model^38^. Further studies are needed to test this hypothesis, and to explore the role of mTORC1-signaling in lipid-induced stress in microglia. Since both mTORC1 activation reduced lipid burden irrespective of APOE levels, and the fact that brain penetrant mTORC1 activators have been identified and are now in clinical development^39,40^, this could represent a therapeutic approach to reduce glial lipid burden.

In conclusion, our study provides an easy and versatile tool for genetic manipulation of iPSC myeloid cells to probe functionality. We demonstrate that it can be deployed in a physiologically relevant arrayed screen for LOAD, and provide insight into regulation of intracellular lipid handling in human microglia.

## Experimental procedures

### iPSC culture and macrophage precursor (preMac) differentiation

Isogenic human iPSC lines (#iPS16, iPS26, iPS36, iPS46) were obtained from Alstembio Inc., and validation iPSC line (BIONi010-C) from Bioneer A/S. All hiPSC lines were maintained in feeder-free conditions in mTeSR plus medium (Stemcell Technologies) on biolaminin 521-coated plates (Biolamina) and were passaged at ∼80% confluence using ReLeSR (Stemcell Technologies) according to manufacturer’s instructions. 10 µM rock inhibitor (Y27632; Callbiochem) was supplied to the medium after thawing for 24 h. All lines were tested for expression of pluripotency markers (TRA1/81, SSEA4; BD Bioscience) and absence of a differentiation marker (SSEA1; BD Bioscience) with flow cytometry prior to each differentiation.

Myeloid factories were generated as previously described (Gutbier et al., 2020; van Wilgenburg et al., 2013). In brief, embryonic bodies (EBs) were formed for 7 days in mTeSR plus medium in AggreWell plates (Stemcell Technologies) with addition of human recombinant BMP-4 (50 ng/ml; R&D), VEGF (50 ng/ml; R&D), and SCF (20 ng/ml; R&D) to induce mesoderm lineage differentiation. Four wells of EBs per line were seeded in a 4-layer cell discs coated with growth factor-reduced matrigel in X-VIVO15 medium (Lonza) supplied with GlutaMAX (Gibco), β mercaptoethanol, penicillin/streptomycin (Gibco), and human recombinant cytokines M-CSF (100 ng/ml; Miltenyi Biotech) and IL-3 (25 ng/ml; Miltenyi Biotech). After six weeks, after preMac viability exceeded 80%, levels of myeloid markers (CD11b, CD16, CD14, CD68; Miltenyi Biotech), and proliferation marker Ki67 (BD Bioscience) were measured on a MACSQuant Analyzer 10 Flow Cytometer (Miltenyi Biotech). Mature preMacs were harvested twice a week for about 3 months for further differentiation and a 50% medium change was carried out with each harvest.

### preMac differentiation into microglia and macrophages

For terminal differentiation into microglia, preMacs were seeded on fibronectin-coated plates (Corning) in DMEM/F12 + GlutaMAX supplied with N2 supplement (Gibco) and containing human recombinant M-CSF (25 ng/ml), IL-34 (100 ng/ml; Miltenyi Biotech), and TGF-β1 (50 ng/ml; PreproTech). 50% of the medium was changed every second day and cells were used for analysis after 10 days of differentiation. Conversely, macrophages were initially cultured without coating in XVIVO15 medium or ImmunoCult for Macrophage Differentiation (Stemcell Technologies) containing 100 ng/ml M-CSF for 6 days, with a 50% medium change on day 3. For nucleofection, macrophages were differentiated in RPMI 1640 with 10% FCS (Gibco) containing M-CSF (100 ng/ml) instead for 6 days (Monkley et al., 2020) (Gutbier et al., 2020) (Wanke et al., 2021).

### sgRNA design (5’-3’orientation)

CD81 sgRNA sequences: UUGGCUUCCUGGGCUGCUA, GCAGCCCUCCACUCCCAUGG, GGCGCUGUCAUGAUGUUCGU

LUC1 (NTC) sgRNA sequences: ACUUACACAUGAGGCGGUA, GUAGUAAAUAUCUAGCUAA, GUCCCUCAGGGUGCAACUU

Sequences for NTC sgRNAs are stated in Table S1. sgRNAs of genes used in arrayed screen and for validation experiments were designed by Synthego (Redwood City, CA). All synthetic sgRNAs were chemically modified with 2′-O-methyl analogs and 3′-phosphorothioate non-hydrolyzable linkages at the first three 5′ and 3′ nucleotides (Synthego, Redwood City, CA).

### Optimized nucleofection protocol for preMacs with subsequent differentiation into microglia

For RNP nucleofection in 96-well format, 25 pmol of sgRNA was complexed with 2 µg of TrueCut Cas9 Protein v2 (Invitrogen) for 20 minutes at room temperature. 50’000 preMacs per reaction were resuspended in 20 µl P3 Primary Cell Nucleofector Solution (Lonza) and mixed with CRISPR/Cas9 RNPs before nucleofection with CM-137 pulse code in a 96-well nucleofection plate using a 4D-Nucleofector 96-well Shuttle nucleofector (Lonza). After nucleofection, cells were allowed to rest for five minutes before 80 µl microglia differentiation medium (see above) was added and the whole volume was transferred to a fibronectin-coated seeding plate (PhenoPlate,

PerkinElmer) containing 100 µl differentiation medium. 75% of the medium was changed the day post-nucleofection, then 50% medium changes were carried out every two days.

### Lipid droplet analysis

Cells were plated in imaging plates (PhenoPlate 96-well, PerkinElmer). PFA-fixed cells were stained with DAPI and 200 nM Nile red (Thermo Fisher) before imaging on an Opera Phenix Plus High-Content Screening System (PerkinElmer). Optical stacks of five images in 1 µm distance were acquired with a 40x water objective in the confocal mode. The 594 nm channel was used to visualize membranes and measure cell morphology parameters such as the cell area with the Harmony Software (version 5.1). Lipid droplets were analyzed using spot segmentation analysis (method B) in the 488 nM channel. Nile red intensity was measured in the 488 nm channel and divided by the cell area.

### Image analysis using Harmony (Perkin Elmer) analysis sequence

Input image: Advanced Flatfield correction and Maximum Projection stack processing

Find nuclei: DAPI channel, analysis using method B, counting objects >20 µm^2^ and setting common threshold to 0.5.

Find cytoplasm: Alexa 568 channel, method A, individual threshold set to 0.06

Morphology properties calculated for region ‘cell’ in population ‘nuclei’ with standard method Intensity properties were calculated in Alexa 488 channel for region ‘cell’ in population ‘nuclei’ Find spots: Alexa 488 channel, ROI nuclei and cell using method B with detection sensitivity 0.47 and splitting sensitivity set to maximum.

### Inhibitor treatment

preMacs were plated at a density of 30’000 cells / well in a 96-well plate (PhenoPlate, PerkinElmer) for microglia differentiated. At day 8 of differentiation, cells were treated with inhibitors for ACAT1 (K604; HY-100400A MedChemExpress, 500 nM), DGAT1 (PF-04620110; HY-13009 MedChemExpress, 1 µM), or mTORC1 (rapamycin, HY-10219 MedChemExpress, 2nM). Cells were fixed with 4% PFA on day 10, and proceeded as above.

### Immunocytochemistry

PFA-fixed cells were blocked with 2% BSA and stained with antibodies against IBA1 (FUJIFILM Wako Pure Chemical Corporation; 019-19741), TMEM119 (Thermo Fisher; PA-62505), or P2YR12 (Sigma; HPA014518) before imaging on an Opera Phenix High Content Imaging System (PerkinElmer).

### Western blot

10 µg of protein was loaded per sample and membrane was probed with antibodies against APOE (Sigma; AB947), RHEB (Abcam; ab25873), TSC2 (Cell Signaling; 4308), phospho-S6 (Cell Signaling; 4858), S6 (Cell Signaling; 2217), and Actin (Abcam; ab49900)

### Flow cytometry

For viability and CD81 expression analysis cells were detached with accutase (Innovative Cell Technologies) and stained with eBioscience Fixable Viability Dye eFluor 780 (Thermo Fisher) and Brilliant Violet anti-human CD45 antibody (Biolegend) for gating and live / dead staining, and PE anti-human CD81 antibody (BD Pharmingen) for expression analysis and were measured on a CytoFLEX LX flow cytometer (Beckman Coulter).

### Cytokine assays

To assess the functional impact of nucleofection and gene KO in iPSC-derived myeloid cells, preMacs were nucleofected with RNPs containing NTC, CD81, or OR52A1 sgRNA, or left untreated (Ctrl). After differentiation into microglia, cells were stimulated with 100 ng/ml LPS or medium for six hours. Supernatant was transferred into a 96-well V-bottom plate and centrifuged for 5 min to remove cellular debris. For cytokine measurements, supernatant was diluted 1:2 with cell culture medium. TNF alpha and IL-6 were quantified using the Procartaplex assays from Invitrogen. To assess functionality of KO in iPSC-derived myeloid cells, macrophages were nucleofected with RNPs containing NTC or TLR4 sgRNAs five days post differentiation. After six days of culture, macrophages were stimulated with 100 ng/ml LPS or 10 µg/ml Poly I:C HMW (Invitrogen) for six hours. Quantification of TNF alpha and IL-6 in cell culture supernatant was performed as described above. For all assays, samples were acquired using a Luminex FlexMap3D system (Thermo Fisher) and analyzed using the BioPlex Software from BioRad.

## Acknowledgements

We would like to thank the Roche Postdoctoral program for funding S.M., and the Roche internships for Scientific Exchange (RiSE) program for funding A.S.G.L. We deeply appreciate critical review and insight of the manuscript from Marcos Costa, Julien Bryois, and Will Macnair.

## Author contributions

Conceptualization S.M., F.R., R.J.,M.K., E.S.M., and L.C.; formal analysis S.M.; investigation S.M, F.W. and A.S.G.L.; methodology S.M. F.W., A.M., and L.T.; project administration S.M.; supervision, M.K., F.R., R.J.; validation S.M. and A.S.G.L.; visualization S.M.; writing – original draft S.M., R.J, F.R. and A.S.G.L.; writing – review & editing S.M., A.S.G.L.,M.K., F.R., and R.J.

