## Supplemental Figures for "An arrayed CRISPR/Cas9 screen identifies mTORC1 as a regulator of lipid droplet accumulation in APOE E3 and APOE KO iPSC-derived microglia"

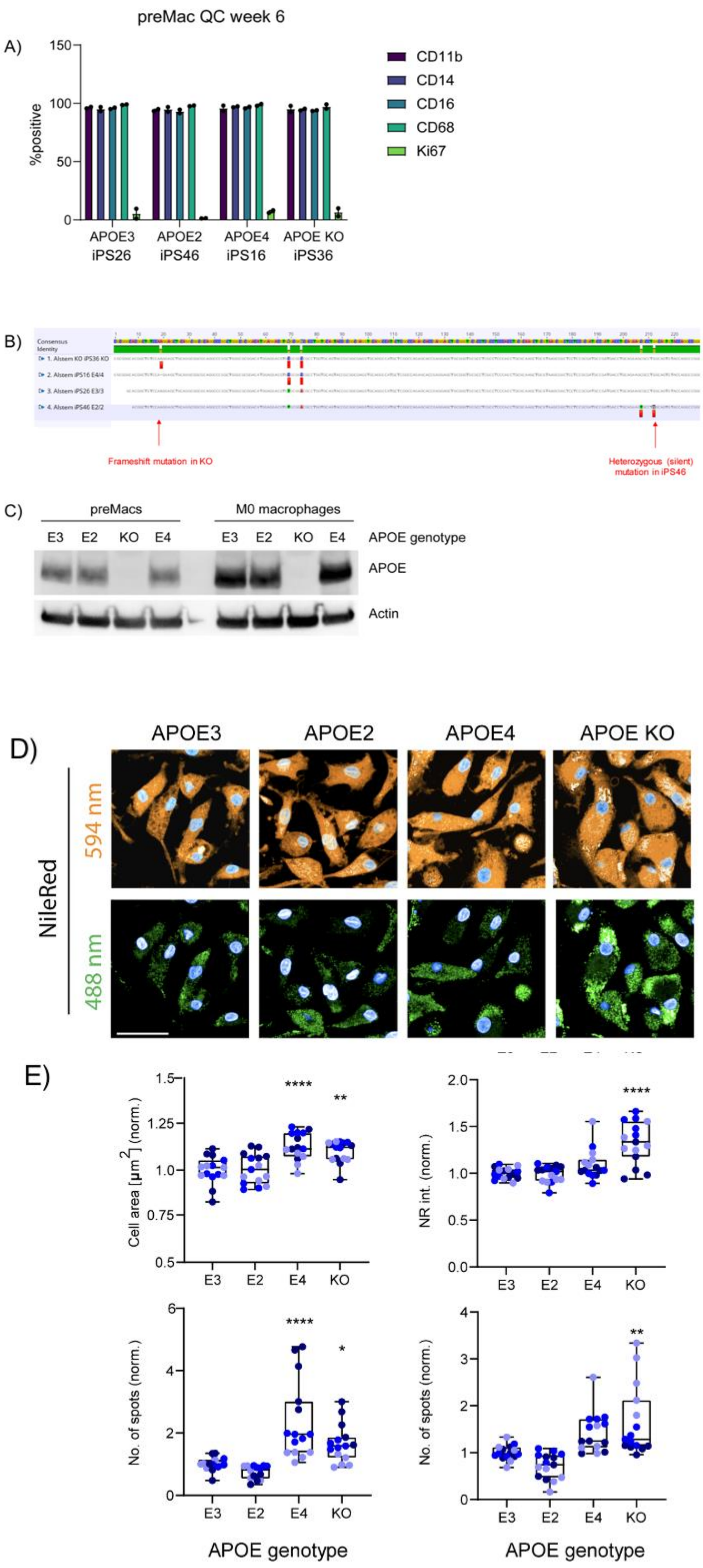
preMac QC week 6
A)
%positive
150
100
50
0
CD11b
CD14
CD16
CD68
Ki67
APOE3 iPS26
APOE2 iPS46
APOE4 iPS16
APOE KO iPS36
B)
Frameshift mutation in KO
Heterozygous (silent) mutation in iPS46
C)
preMacs
M0 macrophages
E3 E2 KO E4 E3 E2 KO E4
APOE genotype
APOE
Actin
D)
APOE3
APOE2
APOE4
APOE KO
NileRed
594 nm
488 nm
E)
Cell area [µm²] (norm.)
NR int. (norm.)
No. of spots (norm.)
No. of spots (norm.)
****
**
****
****
*
**
E3 E2 E4 KO
APOE genotype
APOE genotype

**Figure S1.**

**A)** Flow cytometry analysis of myeloid markers in preMacs harvested six weeks after EB seeding. preMacs were considered mature when >90% of the population tested positive for CD11b, CD14, CD16, and CD68, and <10% for Ki67. Data obtained from two independent differentiations. **B)** Sanger sequencing of isogenic iPSC sets harboring different APOE genotypes. Red markings indicate nucleotides that differ from the APOE3 genotype. **C)** Western blot to confirm APOE expression in isogenic lines at preMac and macrophage stage.

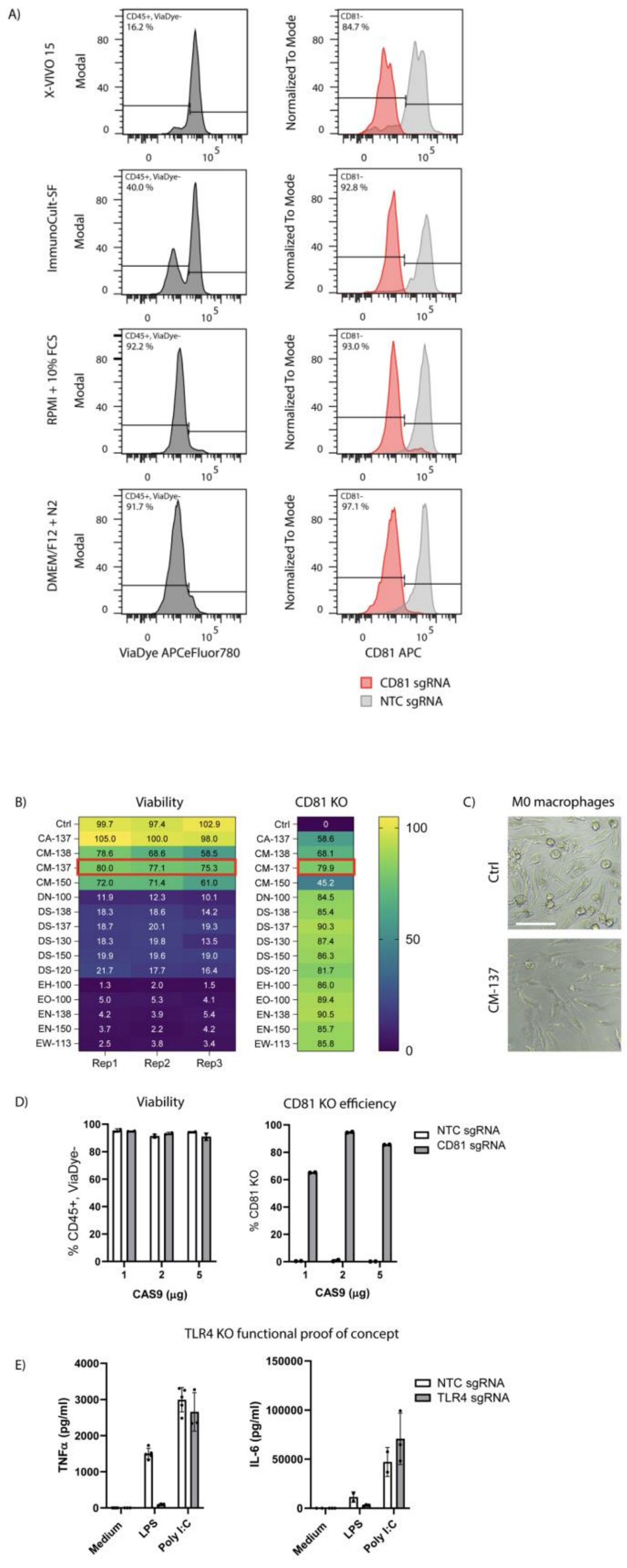

A)
X-VIVO 15
ImmunoCult-SF
RPMI + 10% FCS
DMEM/F12 + N2
Modal
Normalized To Mode
CD45+, ViaDye-
16.2 %
40.0 %
92.2 %
91.7 %
CD81-
84.7 %
92.8 %
93.0 %
97.1 %
ViaDye APCeFluor780
CD81 APC
CD81 sgRNA
NTC sgRNA
B)
Viability
CD81 KO
Ctrl 99.7 97.4 102.9 | 0
CA-137 105.0 100.0 98.0 | 58.6
CM-138 78.6 68.6 58.5 | 68.1
CM-137 80.0 77.1 75.3 | 79.9
CM-150 72.0 71.4 61.0 | 45.2
DN-100 11.9 12.3 10.1 | 84.5
DS-138 18.3 18.6 14.2 | 85.4
DS-137 18.7 20.1 19.3 | 90.3
DS-130 18.3 19.8 13.5 | 87.4
DS-150 19.9 19.6 19.0 | 86.3
DS-120 21.7 17.7 16.4 | 81.7
EH-100 1.3 2.0 1.5 | 86.0
EO-100 5.0 5.3 4.1 | 89.4
EN-138 4.2 3.9 5.4 | 90.5
EN-150 3.7 2.2 4.2 | 85.7
EW-113 2.5 3.8 3.4 | 85.8
Rep1 Rep2 Rep3
C)
M0 macrophages
Ctrl
CM-137
D)
Viability
CD81 KO efficiency
% CD45+, ViaDye-
% CD81 KO
CAS9 (µg)
NTC sgRNA
CD81 sgRNA
TLR4 KO functional proof of concept
E)
TNFα (pg/ml)
IL-6 (pg/ml)
Medium
LPS
Poly I:C
NTC sgRNA
TLR4 sgRNA

**Figure S2: Step-by-step optimization of CRISPR/Cas9 RNP nucleofection**

**A)** iPSC-derived preMacs were differentiated into macrophages for 5 days with M-CSF in following media: X-VIVO15 + 2mM Glutamax + 1% P/S; ImmunoCullt –SF Macrophage medium; RPMI1640 + 2mM Glutamax + 1% P/S + 10% FBS; or DMEM/F12 + N2 supplement. Nucleofection of NTC or CD81 sgRNA with 5 µg CAS9 protein was performed in P3 buffer with 4D Nucleofector (Lonza). Viability and CD81 expression were measured using flow cytometry 5 days post-nucleofection. **B, C)** 5-day macrophages (differentiated in RPMI 1640 + 2mM Glutamax + 1% P/S + 10% FBS) were nucleofected using different pulse codes. Viability and CD81 expression using an antibody against CD81 were measured 5 days post-nucleofection using flow cytometry and CellTiter-Glo Viability Assay (Promega). Rep1-3 indicate technical repeats; wells were pooled to assess CD81 expression. **D)** 25 nmol NTC or CD81 sgRNA was complexed with varying amounts of CAS9 protein before nucleofection using the CM-137 pulse code. Data representative of two independent experiments. **E)** 5-day macrophages were nucleofected with optimized conditions using NTC or TLR4 sgRNA. 6 days post-nucleofection, cells were treated with 100 ng/ml LPS or 10 µg/ml Poly I:C for 6 h and the concentration of TNFα and IL-6 was measured in the supernatant using Luminex assay kit. Data representative of two independent experiments.

A)

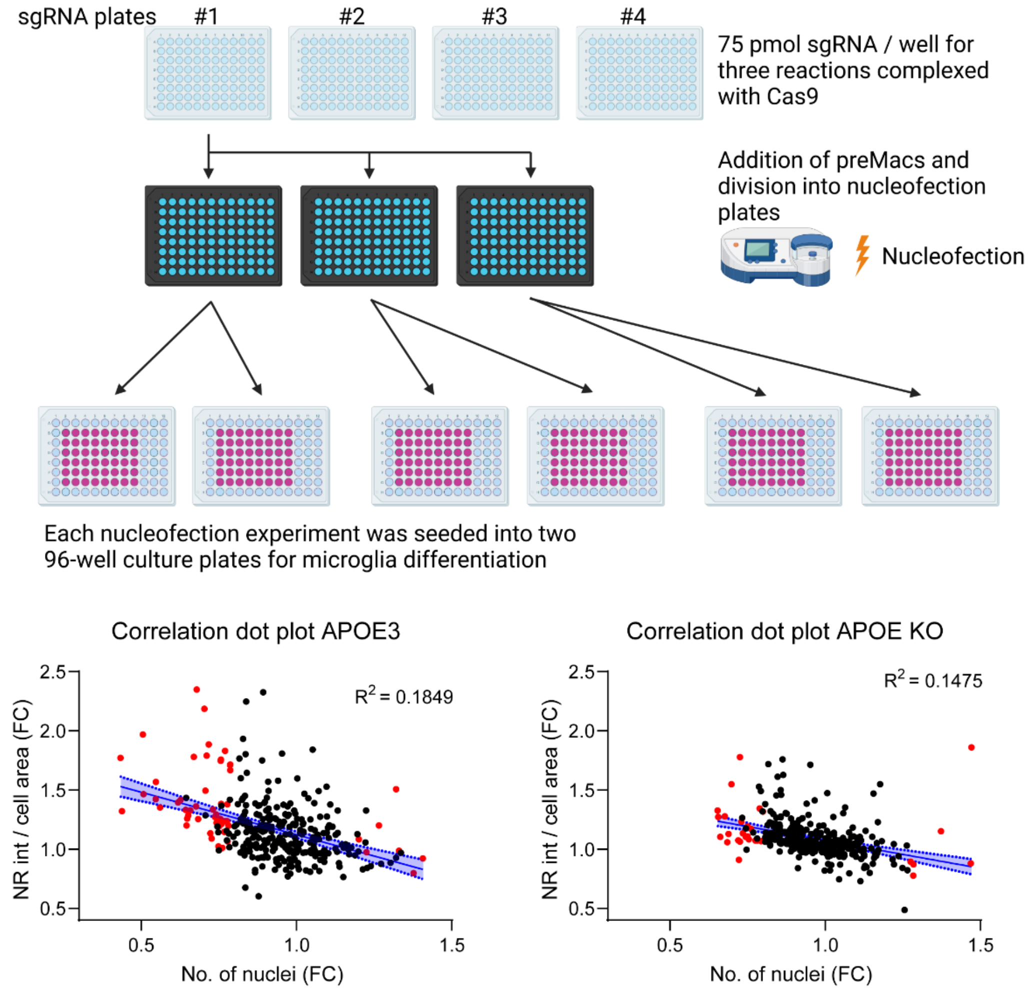


C)

APOE3

| | | |
|---|---|---|
| ABCA8 | FBXO6 | PTCH1 |
| ABCG8 | GNAI1 | PTEN |
| ACAT2 | GRAMD1A | SEMA3B |
| ACOT2 | GRAMD1C | SOAT2 |
| ACSL3 | HMGCR | SREBF1 |
| ANXA13 | HTRA1 | STARD3NL |
| ANXA2 | INSIG2 | TM7SF2 |
| APOC3 | LRP1 | TMEM97 |
| ARL4C | MOV10 | TNFRSF12A |
| ATP13A2 | MVK | VDAC2 |
| ATXN2 | NCOA2 | VDAC1 |
| CREBBP | NPC1 | ARL8B |
| CYP46A1 | NR0B2 | EP300 |
| CYP7A1 | NSDHL | PLAUR |
| EEF2 | OSBPL6 | ACAT1 |
| EEPD1 | OSBPL8 | HSD17B7 |
| ERLIN1 | PLSCR1 | PIP4K2B |
| FABP6 | PPARD | ACSL4 |
| FADS1 | PRKAA1 | |

APOE KO

| | |
|---|---|
| ANXA2 | GPAT3 |
| GRAMD1A | LAMP1 |
| HMGCR | LIPE |
| EHBP1 | SCD |
| SPTLC2 | WSCD1 |
| VPS39 | SULT2B1 |
| HMGCS2 | APOA4 |
| NR0B2 | APOE |
| PNRC1 | RXRB |
| CYP7A1 | S100A11 |
| LRP1 | RAB7A |
| NPC1 | RAN |
| VDAC2 | |
| ACAT2 | |
| FADS1 | |
| PRKAA1 | |
| MED1 | |
| TREM2 | |
| CTSC | |

D)

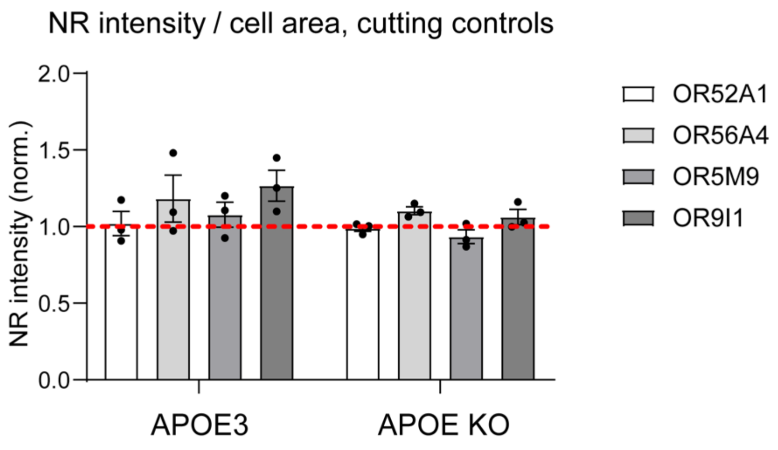


**Figure S3**

**A)** Schematic overview of arrayed screen workflow. 75 pmol target sgRNA / well was complexed with Cas9 and mixed with preMacs in nucleofection plates. Three independent nucleofection experiments were conducted for each sgRNA plate. Wells from each nucleofection plate were seeded in two seeding plates to avoid seeding edge wells, resulting in six plates for analysis of each of the four sgRNA plates. **B)** Correlation plots between the number of nuclei per well and NR intensity / cell area for APOE3 and APOE KO. Linear regression trendline with 95% CI indicates wells with reduced cell number have higher NR intensity reads. Red dots indicate targets with less than 80% or more than 120% cells per well compared to NTC and are significantly different. Significance determined using 1-way ANOVA for each plate. **C)** Gene list of excluded targets for APOE3 and APOE KO lines. **D)** NR intensity per cell area for cutting controls normalized to NTC analyzed by 1-way ANOVA for each line. Results are not significant.

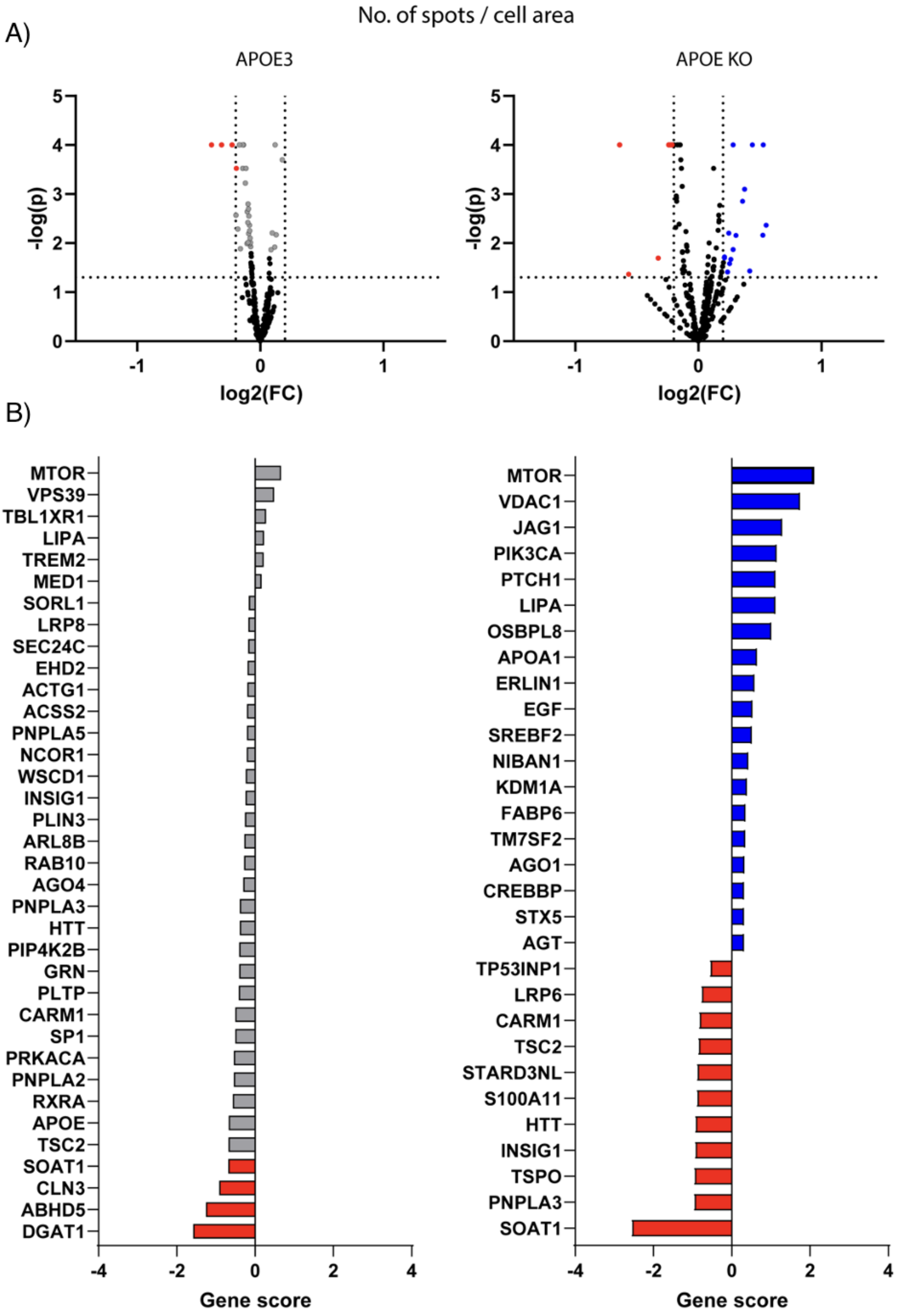
No. of spots / cell area
A)
APOE3
APOE KO
-log(p)
log2(FC)
B)
MTOR
VPS39
TBL1XR1
LIPA
TREM2
MED1
SORL1
LRP8
SEC24C
EHD2
ACTG1
ACSS2
PNPLA5
NCOR1
WSCD1
INSIG1
PLIN3
ARL8B
RAB10
AGO4
PNPLA3
HTT
PIP4K2B
GRN
PLTP
CARM1
SP1
PRKACA
PNPLA2
RXRA
APOE
TSC2
SOAT1
CLN3
ABHD5
DGAT1
Gene score
MTOR
VDAC1
JAG1
PIK3CA
PTCH1
LIPA
OSBPL8
APOA1
ERLIN1
EGF
SREBF2
NIBAN1
KDM1A
FABP6
TM7SF2
AGO1
CREBBP
STX5
AGT
TP53INP1
LRP6
CARM1
TSC2
STARD3NL
S100A11
HTT
INSIG1
TSPO
PNPLA3
SOAT1
Gene score

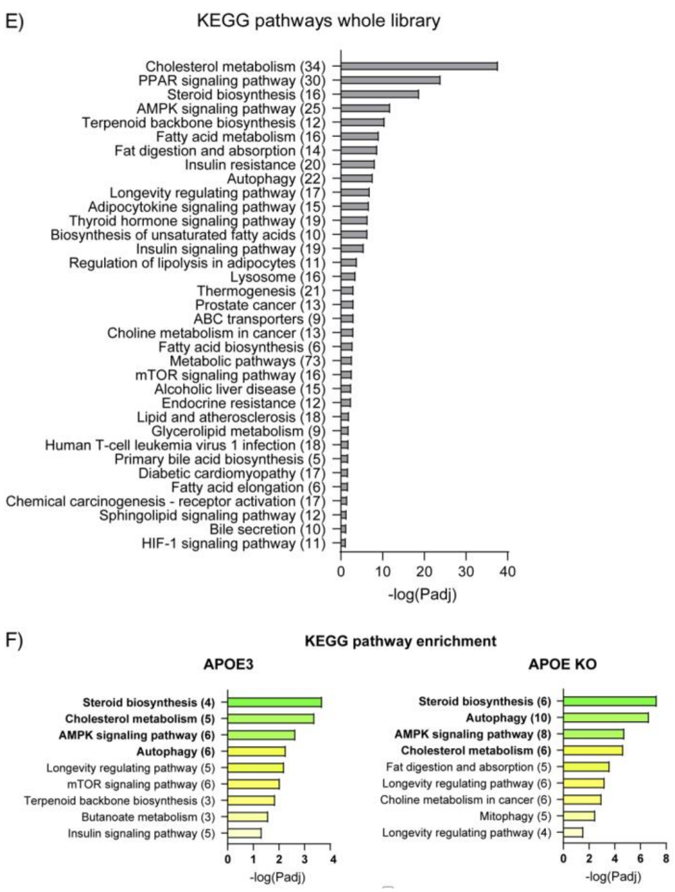


**Figure S4**

**A, B)** Volcano plots and gene scores indicating significant hits for APOE3 and APOE KO for no. of spots per cell. Red and blue indicate significant hits with a fold change of at least 15%, grey indicate significant hits in APOE3 that did not make the 15% fold change cut-off. Data combined from three nucleofection experiments per gene. All genes analyzed separately for each experiment with paired 1-way ANOVA and comparison to NTC on each plate. Gene score = log2(FC) * -log(p). **C)** KEGG pathway analysis of the whole focused library, 334 genes. **D)** KEGG pathway analysis of significant screen hits for both APOE3 and APOE KO. All pathway analyses were performed using g:Profiler (version e109_eg56_p17_1d3191d)).
